# Hijacking the transcriptional activation potential of the BAF complex via Induced Proximity

**DOI:** 10.1101/2023.12.18.572217

**Authors:** Lionel Chia, David Mayhew, Brenna Sherbanee, David L. Lahr, Asad M. Taherbhoy

**Author notes:** Corresponding Author: Asad M. Taherbhoy.

## Abstract

The BAF (Brg/Brahma-associated factors) complex, also referred to as the mammalian Switch/Sucrose-Nonfermentable (mSWI/SNF) chromatin remodeling complex, plays a pivotal role in epigenetically regulating diverse transcriptional programs. BAF’s chromatin remodeling activity, which enhances accessibility to transcriptional machinery, is critical for gene regulation. Recent studies have demonstrated that redirecting BAF complexes to bivalent promoters can alter the local epigenetic landscape, creating a permissive environment for transcription. As such, we hypothesize that redirecting BAF to “turn on” therapeutically relevant genes offers a potential approach for disease treatment.

Using rapamycin as a chemical inducer of proximity (CIP) via CRISPR/Cas9 and FKBP/FRB dimerization, we redirected BAF complexes to the promoter of fetal hemoglobin (*HBG*), a therapeutic target for beta-globinopathies like sickle cell anemia and beta-thalassemia. This resulted in changes to the local chromatin and epigenetic landscapes, and increased *HBG1* expression. Having confirmed BAF’s ability to activate gene expression, we then performed a genome-wide CRISPR activation drop-out screen to identify genes that when activated by BAF, suppress cellular proliferation. In addition to known tumor suppressors, our screen identified a number of genes with the ability to inhibit cell proliferation when activated by BAF. Collectively, our findings highlight the potential for harnessing the BAF’s intrinsic transcriptional activation capabilities for therapeutic purposes and lays the foundation for the potential development of therapeutics that function via induced proximity.

## Introduction

The BAF (BRG1/BRM-associated factor) chromatin remodeling complex, also known as the mSWI/SNF complex, is a highly conserved ATP-dependent multi-subunit protein that regulates transcription and governs genome architecture^1–3^. The BAF complex subunits, which are encoded by 29 genes, assemble into three types of BAF complexes: the canonical BAF complex (cBAF), the non-canonical BAF complex (ncBAF), and the polybromo-associated BAF complex (PBAF)^4–6^. Each of these complexes contains not only common and complex specific subunits but also the mutually exclusive ATPase domain-containing subunit BRG1 or BRM^7–9^. This enables the BAF complex to utilize energy from ATP hydrolysis to remodel chromatin via nucleosome sliding and eviction^10–13^. Furthermore, its interactions with transcription factors and co-regulators, coupled with its binding to various genomic elements (enhancers, promoters, and CTCF binding sites), allow for precise transcriptional control, making it essential for a variety of physiological processes such as cell differentiation, development, and DNA repair^1,2,11,14–17^. Dysregulated BAF function arising from mutations within BAF subunits accounts for more than 20% of all cancers^9,18,19^, resulting in altered gene expression patterns and tumorigenesis^9,19,20^. As a result, the BAF complex has emerged as an important therapeutic target in cancer.

In prior studies, *in vitro* overexpression of various BAF subunits changes its global transcriptomic and chromatin profile, highlighting its transcriptional potential^21–23^. Overexpression of *BAF47*, a core BAF complex subunit, in BAF47 deficient malignant rhabdoid tumor (MRT) cell lines increase genome-wide chromatin accessibility, global enhancer activation, and expression of genes with bivalent promoters (defined by the presence of H3K4me3 and H3K27me3 marks)^21,24^. Similarly, overexpression of *BRG1* increases global chromatin accessibility and activates genes with tumor-suppressive properties in both lung cancer and small cell carcinoma of the ovary, hypercalcemic type (SCCOHT)^22,23^. Of particular note, using chemical inducers of proximity (CIP) such as rapamycin to redirect BAF to various genomic loci, increases chromatin accessibility and expression of bivalent genes in mouse embryonic stem cells^25,26^. This suggests that hijacking BAF to activate genes transcriptionally could serve as a novel therapeutic opportunity.

In recent years, induced proximity-based drugs have garnered attention and revolutionized drug development. In general, induced proximity drugs can change the fate of a target protein or modulate various biological processes by bringing the target and effector proteins close together^27^. Proteolysis-targeting chimeras (PROTACs) and molecular glue degraders are examples of proximity-based drugs that bring a target protein to close proximity to a ubiquitin E3 ligase, causing target protein ubiquitination and degradation^27^. Beyond targeted protein degradation, small molecule-induced proximity modalities have been used to induce post-translational modifications (e.g. phosphorylation), protein stabilization (de-ubiquitination), and rewire cellular function (e.g., CAR-T signaling and function)^27,28^. Here, we aim to understand the transcription activation potential of the BAF complex by inducing its proximity to target genes.

To explore the therapeutic potential of hijacking BAF and its inherent transcriptional activity, we established the FKBP/FRB inducible recruitment for epigenome editing by Cas9 (FIRE–Cas9) system in various cellular models^25^. This allows the BAF complex to be recruited to any genomic region of interest via CIP using rapamycin. Using fetal hemoglobin (HBG) as a model system, we show that targeted recruitment of the BAF complex to the promoter increases local chromatin accessibility, enriching for active histone marks while depleting repressive epigenetic marks along the *HBG* genomic loci, driving *HBG* expression at the mRNA and protein levels. By performing a CRISPR-based screen in K562 cells, we further demonstrate the robustness of BAF’s inherent ability to activate genes that have the potential to suppress cellular proliferation. Interestingly, genes that BAF activated extended beyond known tumor-suppressor genes, activating additional genes that could be potential oncology targets for the induced proximity approach. Our findings suggest that we can leverage BAF’s transcriptional activity as a novel modality to treat diseases.

## Results

### BAF complexes can be directed to genomic loci of interest to increase chromatin accessibility

To assess the transcriptional potential of the BAF complex, we first stably expressed the FIRE-dCas9 system^25^ in two cell-based models, HEK293T (293T-FIRE) cells containing the SV40 large T-antigen and the chronic myelogenous leukemia K562 cells (K562-FIRE)^29,30^. Both these lines express BAF with no known damaging mutations^31^. The ectopically expressed FIRE-dCas9 system consists of a catalytically dead Cas9, an MS2 scaffold protein fused to a FKBP (FK506 binding protein) domain, and the SS18 subunit fused to the FRB domain at the N-terminus^25^ (**Supplemental Figure 1A-B**). The SS18 fusion protein allows for incorporation into a functional BAF unit^26^ (**Supplemental Figure 1C-D**), and, combined with specific sgRNAs and rapamycin as a CIP, allows the BAF complex to specifically target genomic regions of interest^25^ **(Supplemental Figure 1A)**.

As BAF has been shown to be able to activate a variety of genes with rapid and robust gene expression at specific developmentally primed bivalent promoters^21,22,25^, we wanted to determine if hijacking BAF could activate genes that have potential therapeutic value. To this end, we used fetal hemoglobin as a model system as globin genes have few downstream targets that could confound our results^32,33^. Furthermore, its overexpression serves as a potential therapeutic opportunity to treat beta-globinopathies such as sickle cell anemia and beta-thalassemia^34–36^.

To test the ability of BAF to activate HBG expression, we started by stably expressing pooled small guide RNAs (sgRNA) targeting the *HBG1* promoter (sg*HBG1* 1: −45; sg*HBG1* 2: −120; and sg*HBG1* 3: −183 bp upstream of the transcriptional start site). To determine if the BAF complex is being recruited to the promoter, we performed a chromatin-immunoprecipitation (ChIP) across the *HBG1* loci (**Figure 1A**) targeting the V5-tagged SS18-FRB and BRG1. Indeed, upon rapamycin treatment, the BAF complex was recruited to the *HBG1* promoter at sites of sg*HBG1* binding in K562-FIRE and 293T-FIRE cells (**Figure 1B and 1C**). Non-targeting sgRNA (sgNT) serves as a control to eliminate the effects of rapamycin. Of the three distinct BAF complexes, cBAF, PBAF, and ncBAF^4–6^, only cBAF and ncBAF contain the SS18 subunit. We wanted to determine which BAF complexes were recruited to the *HBG1* loci. Using ARID1a and GLTSCR1 to identify cBAF and ncBAF complexes, respectively, we observed an enrichment of ARID1a and GLTSCR1 binding over sg*HBG1* targeted sites in both K562-FIRE and 293T-FIRE cells following rapamycin treatment (**Figure 1B and 1C**). As expected, ARID2, unique to PBAF, is not recruited to the *HBG1* loci (**Supplemental Figure 1E-F**). Having confirmed the recruitment of BAF to the *HBG1* loci, we next assessed the ability of BAF to “open” chromatin using an assay for transposase assessable chromatin (ATAC). In both K562-FIRE and 293T-FIRE cells, recruitment of BAF to the *HBG1* loci increases local chromatin accessibility (**Figure 1D-E**). Taken together, our cellular system allows for the recruitment of BAF to our genomic site of interest, which in turn is sufficient to open chromatin in that region.

**Figure 1.**
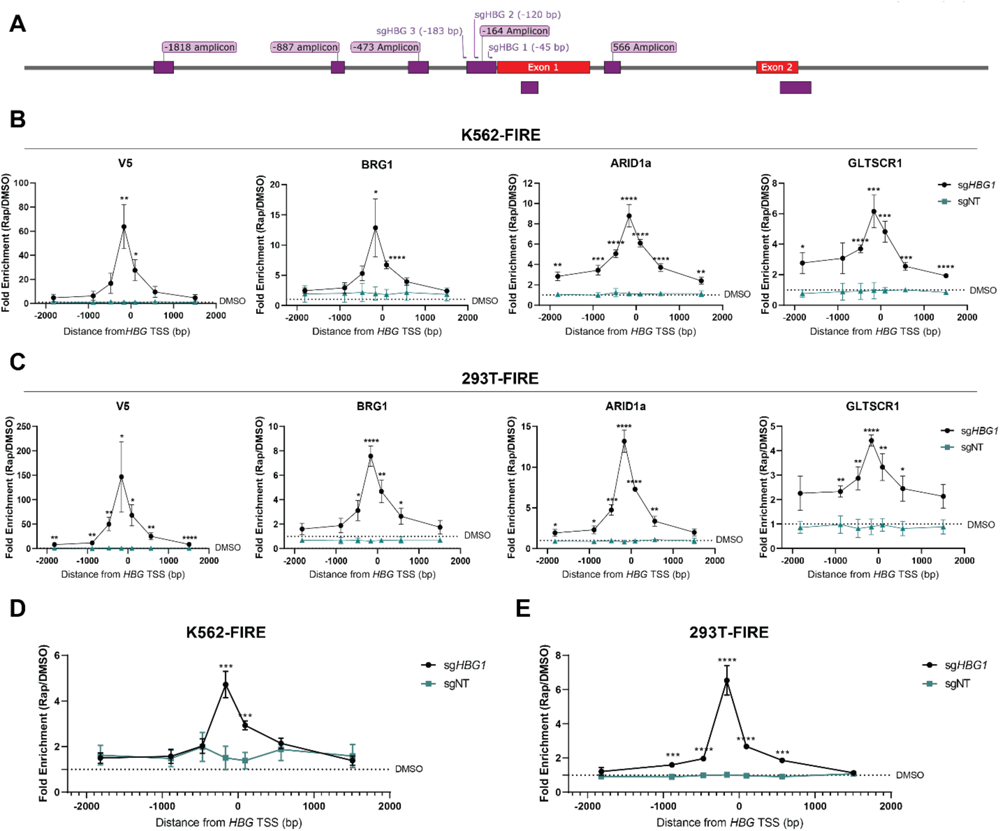
Targeted cBAF and ncBAF complex recruitment to the genomic region of interest increases chromatin accessibility. **(A)** A Schematic representation of the hemoglobin gamma 1 (*HBG1*) loci showing the locations of sgRNA binding and regions for qPCR analysis. **(B)** ChIP-qPCR comparing BAF complex occupancy across the *HBG1* loci in K562-FIRE cells expressing sg*HBG1* or sgNT after 24 h of 10 nM rapamycin (Rap) or DMSO treatment. **(C)** ChIP-qPCR comparing BAF complex occupancy across the *HBG1* loci 293T-FIRE cells expressing sg*HBG1* or sgNT after 24 h of 10 nM rapamycin or DMSO treatment. **(D)** ATAC-qPCR of comparing chromatin accessibility of K562-FIRE cells expressing either sg*HBG1* or sgNT after 24 h of 10 nM rapamycin or DMSO treatment. **(E)** ATAC-qPCR of comparing chromatin accessibility of 293T-FIRE cells expressing either sg*HBG1* or sgNT after 24 h of 10 nM rapamycin or DMSO treatment. All data are shown from 3 independent experiments performed in triplicate. Data presented as mean ± SEM. Significance was determined by 1-way ANOVA with Turkey’s multiple-comparisons test, with indicated significance showing the comparisons between sg*HBG1* and sgNT. \**P* < 0.05, \*\**P* < 0.01, \*\*\**P* < 0.001, \*\*\*\**P* < 0.0001.

### Recruitment of BAF to the *HBG1* promoter induces gene and protein expression

Having established that BAF can be redirected to genomic sites of interest to open chromatin, we next investigated if this would lead to fetal hemoglobin expression at both the mRNA and protein levels. Interestingly, following rapamycin induction, levels of *HBG1* mRNA were significantly increased in K562-FIRE cells (**Figure 2A**). In addition to increased mRNA levels, we could also detect increased HBG protein expression (**Figure 2B**). Of note, endogenous levels of *HBG1* appear to be lower in K562-FIRE cells with sg*HBG1* than in cells with sgNT (**Figure 2B**). It has been observed that dCas9 recruitment to sgRNA sites on promoters sterically hinders transcription factors from accessing the promoter to drive gene expression, decreasing endogenous transcription levels^37^. Despite that, BAF could still increase *HBG1* expression when recruited to the *HBG1* loci, giving credence to its transcription activation potential. Similarly, in 293T-FIRE cells, BAF was able to increase *HBG1* mRNA expression robustly (**Figure 2C**). Because endogenous *HBG1* levels in 293T-FIRE cells are low, detecting increased HBG protein levels proved challenging. To this end, we used the highly sensitive fluorescence-activated cell sorting (FACS) assay to detect increases in intracellular HBG upon BAF recruitment. Albeit subtle, we detected increased HBG levels in 293T-FIRE upon recruitment of BAF to the *HBG1* loci (**Figure 2D-2E**). A difference in overall HBG expression in K562-FIRE and 293T-FIRE highlights the importance of the native cellular context and specific transcription factors or co-factors that may be required to act in concert with the BAF complex to activate transcription. Nevertheless, our data demonstrates that recruitment of BAF to the *HBG1* loci can increase both HBG1 transcript and protein levels.

**Figure 2.**
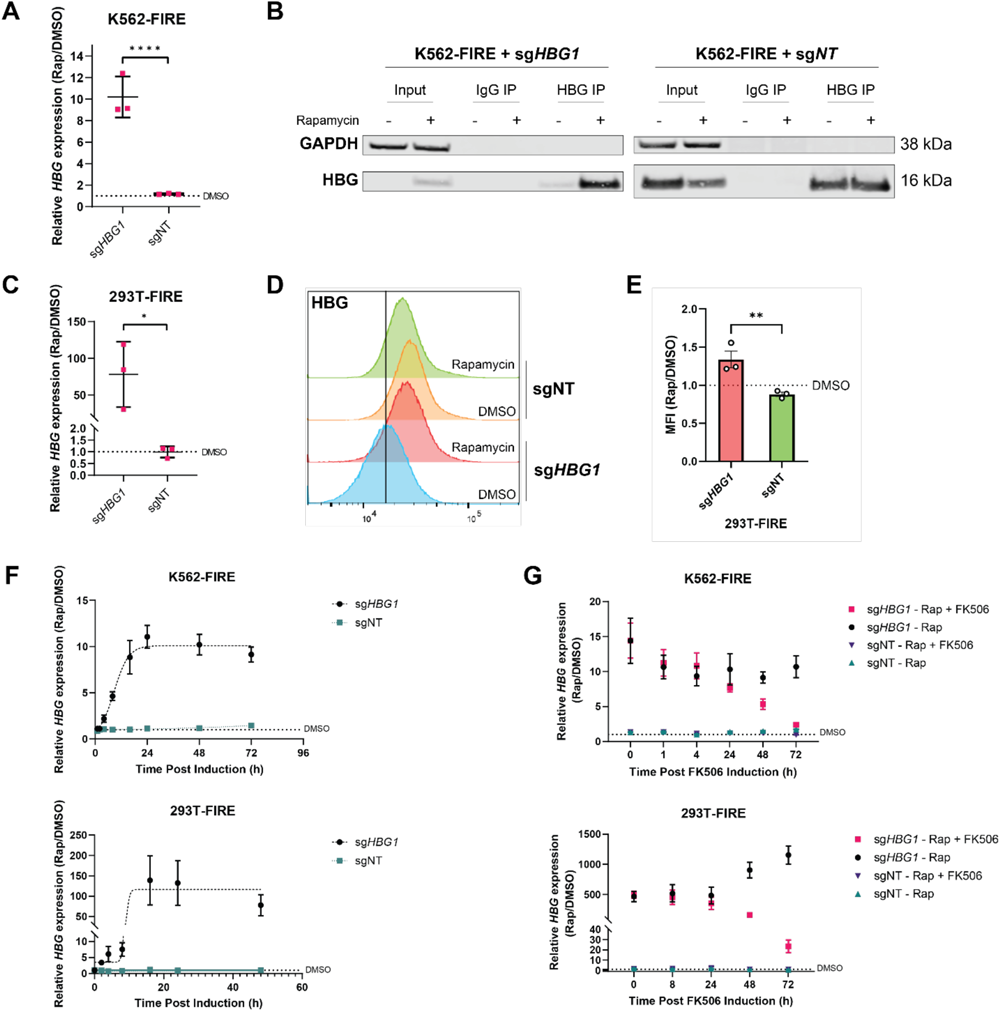
Targeted recruitment of BAF to the *HBG1* promoter induces gene expression at the mRNA and protein levels. **(A)** *HBG1* expression in K562-FIRE cells expressing sg*HBG1* or sgNT after 48 h of 10 nM rapamycin or DMSO treatment. **(B)** Representative immunoblot of fetal hemoglobin (HBG) after immunoprecipitation in K562-FIRE expressing sg*HBG1* or sgNT after 72h of 10 nM rapamycin or DMSO treatment. **(C)** *HBG1* expression in 293T-FIRE cells expressing sg*HBG1* or sgNT after 48h of 10 nM rapamycin or DMSO treatment. **(D)** Representative flow cytometric profile of *HBG1* in 293T-FIRE cells expressing sg*HBG1* or sgNT after 72 h of 10 nM rapamycin or DMSO treatment. **(E)** Comparison of median fluorescence intensities (MFI) of *HBG1* protein levels in 293T-FIRE cells expressing sg*HBG1* or sgNT after 72 h of 10 nM rapamycin or DMSO treatment. **(F)** *HBG1* expression in K562-FIRE (upper panel) and 293T-FIRE (lower panel) cells expressing sg*HBG1* or sgNT after 10 nM of rapamycin or DMSO treatment across various time points. **(G)** *HBG1* expression when FK506, a rapamycin competitor, is added to rapamycin-treated K562-FIRE (upper panel) and 293T-FIRE (lower panel) cells expressing sg*HBG1* or sgNT. 10 nM of FK506 or DMSO was added to cells after 24 h of 10 nM rapamycin treatment. 10 nM rapamycin was maintained throughout the experiment. Relative *HBG1* expression is normalized to DMSO-treated cells. All data shown are from 3 independent experiments with qPCR experiments performed in triplicates. Data presented as mean ± SEM. Significance was determined by 1-way ANOVA with Turkey’s multiple-comparisons test, with indicated significance showing the comparisons between sg*HBG1* and sgNT. \**P* < 0.05, \*\**P* < 0.01, \*\*\*\**P* < 0.0001.

To understand the kinetics of transcriptional activation by BAF, we assessed *HBG1* expression across different time points. Remarkably, BAF-mediated gene expression was rapid, with half-maximal *HBG1* expression detected at 2.4 h and 9.4 h in K562-FIRE and 293T-FIRE cells, respectively (**Figure 2F**). Maximum gene expression was observed by 24 h in both K562-FIRE and 293T-FIRE cells (**Figure 2F**). To determine if the induced *HBG1* expression was reversible, FK506 was added to compete with the rapamycin-FKBP interaction, reversing BAF’s recruitment to the *HBG1* promoter. *HBG1* expression notably starts decreasing 24 h after FK506 addition, with *HBG1* expression reverting to endogenous levels at about 72 h post FK506 induction in both K562-FIRE and 293T-FIRE cells (**Figure 2G**). This confirms that the transcriptional activity of BAF can be rapidly induced and reversed, as previously reported^25^.

Based on global accessibility assays (ATAC-seq) and ChIP-sequencing, the BAF complex regulates gene expression in part by remodeling chromatin around the enhancers^21,38–40^. We sought to determine if BAF could also mediate *HBG1* gene expression when redirected to the enhancer regions of *HBG1*. We designed sgRNAs to direct the BAF complex to enhancer regions denoted by DNase I hypersensitivity (HS1-HS4) within the locus control region of the hemoglobin loci. These enhancer regions play a crucial role in the developmental control of hemoglobin expression, and it was previously shown that activating these enhancer regions via P300 recruitment increases the expression of globin genes such as embryonic hemoglobin (*HBE*), and beta-hemoglobin (*HBB*)^41–43^. However, recruiting BAF to the enhancer regions did not significantly increase the expression of globin genes *HBG1, HBE* and *HBB* in either cell line (**Supplemental figure 2A-B**). In addition, further ChIP-sequencing analysis of Histone 3 lysine 4 monomethylation (H3K4me1) and Histone 3 lysine 27 acetylation (H3K27Ac), coupled with ATAC-sequencing and genomic perturbation studies have identified a locus within the *BGLT3* gene and a locus 4-kilo base pairs upstream of the *HBG* promoter (HBG-4KBP) to be novel enhancers that regulate *HBG1* expression^44–46^. We therefore designed sgRNAs to recruit BAF to these novel loci. Here again, when BAF was recruited to these novel enhancers, no significant expression of globin genes was observed in either cell line (**Supplemental Figure 2C-D**). Taken together, recruitment of the BAF complex to the *HBG1* promoter, but not the enhancer region, induces *HBG1* expression rapidly and reversibly.

### BAF complexes directed to loci of interest cause local epigenetic changes

As the BAF complex can bind and modify local epigenetic states^25,26^, we wanted to understand if recruiting BAF to the *HBG1* loci was accompanied by changes in epigenetic marks. After 24 h of rapamycin induction, we observed the enrichment of trimethylation at Histone 3, lysine 4 (H3K4me3), a marker for active promoters^47^, within the gene body in K562-FIRE cells (**Figure 3A**). Surprisingly in 293T-FIRE cells, rather than the enrichment of H3K4me3 marks, it was H3K27Ac that was enriched over sites of BAF recruitment (**Figure 3B**). H3K27Ac epigenetic marks are typically associated with active enhancers, but when located near promoters, they play a role in transcriptional elongation^48–50^.

**Figure 3.**
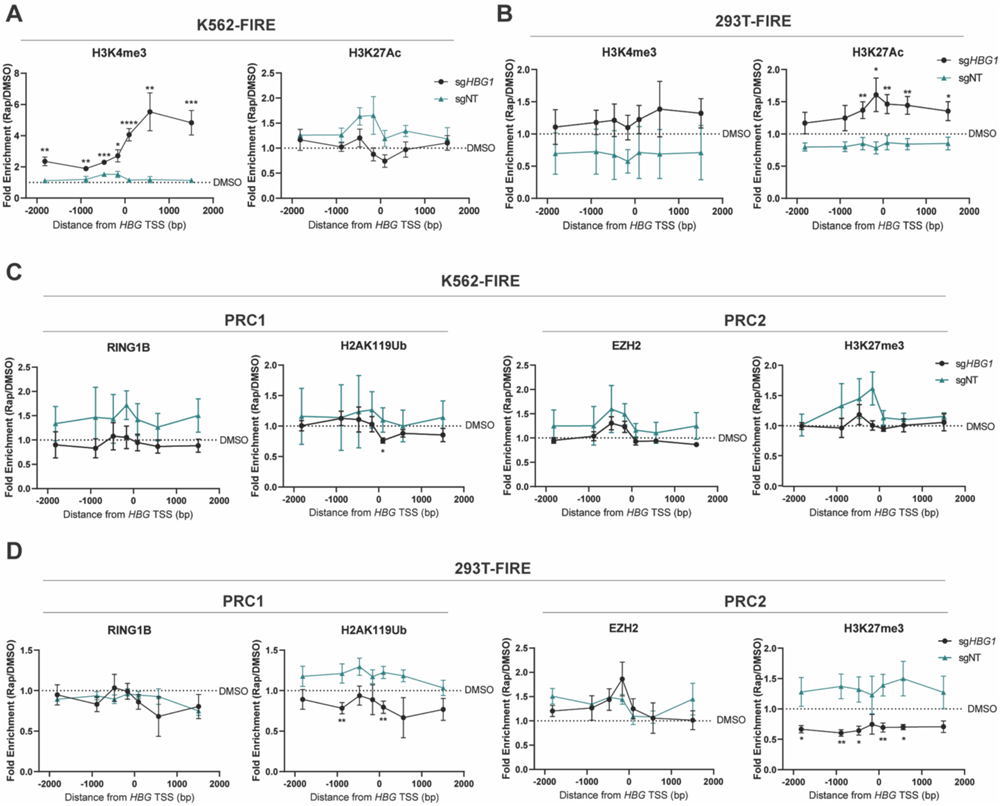
Targeted BAF-mediated gene activation changes local epigenetic states. **(A)** ChIP-qPCR for H3K4me3 and K3K27Ac active histone marks on the *HBG1* loci in K562-FIRE cells expressing sg*HBG1* or sgNT after 24 h of 10 nM rapamycin or DMSO treatment. **(B)** ChIP-qPCR for H3K4me3 and K3K27Ac active histone marks on the *HBG1* loci in 293T-FIRE cells expressing sg*HBG1* or sgNT after 24 h of 10 nM rapamycin or DMSO treatment. **(C)** ChIP-qPCR for repressive histone marks of the PRC1/2 complexes on the *HBG1* loci in K562-FIRE cells expressing sg*HBG1* or sgNT after 24 h of 10 nM rapamycin or DMSO treatment. **(D)** ChIP-qPCR for repressive histone marks of the PRC1/2 complexes on the *HBG1* loci in 293T-FIRE cells with either sg*HBG1* or sgNT after 24 h of 10 nM rapamycin or DMSO treatment. Data are presented as mean ± SEM from 3 independent experiments performed in triplicates. Significance was determined by 1-way ANOVA with Turkey’s multiple-comparisons test, comparing sg*HBG1* with sgNT. \**P* < 0.05, \*\**P* < 0.01, \*\*\**P* < 0.001, \*\*\*\**P* < 0.0001.

To create a more permissive environment for transcription, BAF opposes heterochromatin formation and remodels chromatin to create a more “open” euchromatin state^26,51^. We next wanted to determine if targeted BAF recruitment affected heterochromatin status by monitoring post-translational changes mediated by the Polycomb repressive complex 1 and 2 (PRC1 and 2) at the *HBG1* loci. In K562-FIRE cells, no significant changes were observed across the *HBG1* region except a slight decrease in the enrichment of monoubiquitinated Histone 2A at lysine 119 (H2AK119Ub) at the gene body (**Figure 3C**). H2AK119Ub is associated with gene repression and is essential in PRC1-mediated transcription repression^52^. Of note, as *HBG1* baseline expression is high (**Figure 2B**) in K562 cells, the effects of H2AK119Ub loss could be negligible.

On the contrary, in 293T-FIRE cells, which simulate a “normal” model, low *HBG1* suggests the presence of heterochromatin states and transcriptional silencing programs^53–55^. Indeed, we observed that BAF recruitment not only increases chromatin accessibility (**Figure 1E**) but induces a loss of H2AK119ub and trimethylation at Histone 3, lysine 27 (H3K27me3) across the *HBG1* loci (**Figure 3D**); H3K27me3 is an epigenetic mark associated with gene silencing^56^. Although differences are observed across cell lines, targeted BAF recruitment is associated with the enrichment of active epigenetic marks and depletion of repressive epigenetic marks.

### BAF activates genes with various epigenetic states and basal expression levels

Having shown the ability of BAF to activate *HBG1*, we wanted to explore the potential of BAF to activate genes across the genome in an unbiased way that could potentially be leveraged from a therapeutic perspective. To this end, we performed a genome-wide CRISPR screen to identify genes that, when activated by BAF, would result in cell death or suppressed proliferation (**Figure 4A**). We first established a pool of K562-FIRE cells stably expressing sgRNAs from the Synergistic Activation Mediator (SAM) sgRNA library^32^. The SAM library consists of 70,290 MS2-modified sgRNAs that target 23,430 unique protein-coding genes in the human genome, with an average coverage of three sgRNAs per gene^32^. These pooled cells were selected for at least 14 days to eliminate any detrimental cellular effects due to the inhibition of gene expression when dCas9 localizes to promoters. These pooled cells were subsequently treated with either DMSO or rapamycin to induce BAF recruitment to promoters (**Figure 4A**).

**Figure 4.**
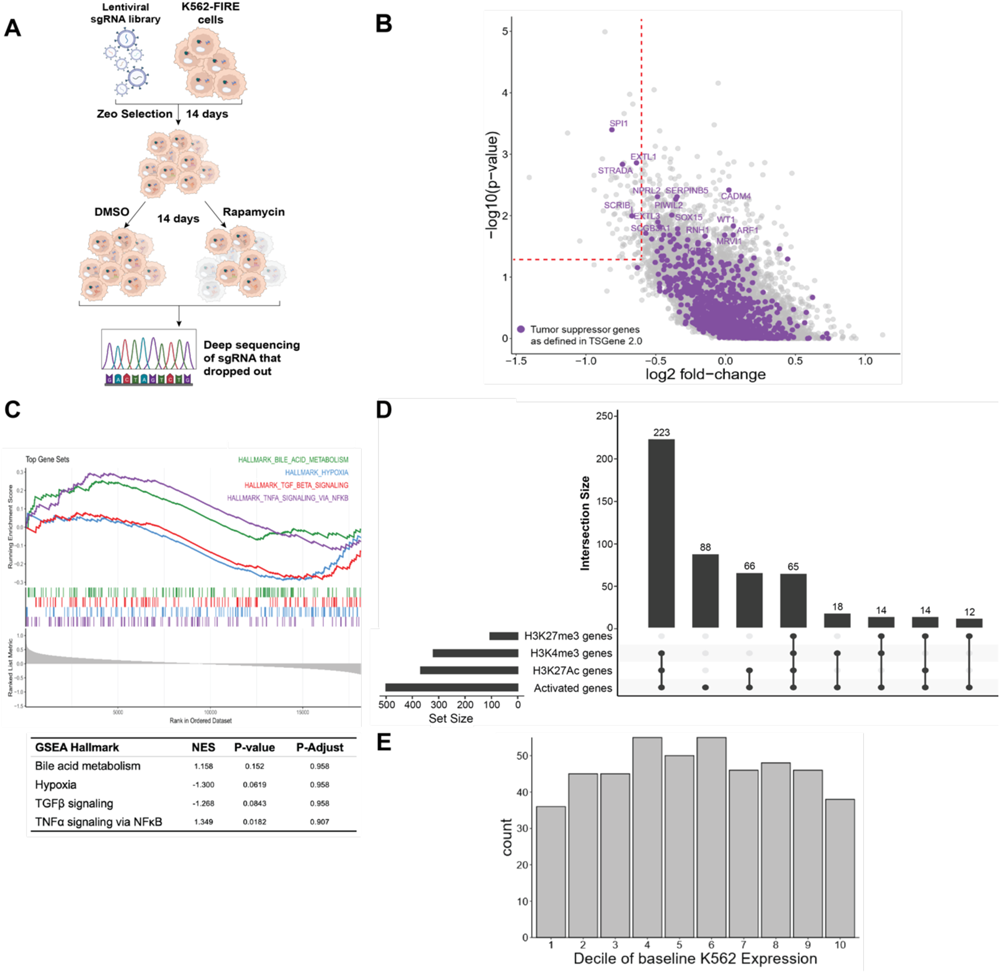
BAF-mediated gene expression is robust and has therapeutic potential. **(A)** Schema of whole genome-wide CRISPR screen to identify genes that suppress cell proliferation when activated through recruitment of endogenous BAF. **(B)** Volcano plot of BAF-activated genes that inhibit cell proliferation analyzed using MAGeCK. Tumor suppressors are indicated in purple, and grey dots indicate other genes that have the propensity to inhibit cell proliferation. Dashed lines indicate thresholds of *P* < 0.05 and Log_2_[Fold Change] < −0.6. Data shown are from 3 independent sgRNA library transductions. **(C)** GSEA plots of the most significant Hallmark gene sets for BAF activated genes. **(D)** Epigenetic marks at the promoter of the top 500 genes that BAF activates. **(E)** Baseline RNA expression of the top 500 genes activated by BAF binned in 10% increments, with bin 1 indicating genes with low RNA expression and bin 10 being the highest RNA expression.

Because rapamycin has pharmacological effects on cells, we first sought to determine if we could normalize the effect of rapamycin on gene expression with K562-FIRE cells expressing sg*HBG1* or sgNT. Principal component analysis of the whole transcriptome separates DMSO control or rapamycin-treated into distinct groups, with rapamycin-treated groups being more similar to each other than the DMSO control-treated groups (**Supplemental Figure 3A**). In K562-FIRE cells, *HBG1/2* has a high endogenous expression level, which increases when BAF is recruited to its promoter (**Supplemental Figure 3B-D**). BAF-mediated gene activation enriches *HBG2*, which is also targeted by sg*HBG1* due to its high DNA sequence similarity. (**Supplemental figure 3B-D**). RNA-sequencing confirms dCas9’s suppressive activity in sg*HBG1*, and despite a significant fold difference in the sg*HBG1* group upon rapamycin treatment, BAF-mediated *HBG1* expression did not reach NT levels (**Supplemental Figure 3C**).

The differentially expressed genes between rapamycin and DMSO in K562-FIRE cells expressing sg*HBG1* included 690 upregulated genes and 1,075 downregulated genes, with strong upregulation of *HBG1* and *HBG2* observed (**Supplemental Figure 3D**). Similarly, in K562-FIRE cells containing sgNT, rapamycin, and DMSO differentially expressed genes included 67 upregulated genes and 175 downregulated genes (**Supplemental Figure 3E**). When comparing rapamycin-treated K562-FIRE cells containing sg*HBG1* with rapamycin treated cells containing sgNT, only three genes were upregulated and downregulated (**Supplemental Figure 3F**), with all differentially expressed genes located within the hemoglobin loci (*BGLT3* and *HBBP1* are non-coding regulator genes^45^). However, it is important to note that the reduced number of statistically significant genes in these comparisons is partly due to the reduced number of replicate samples in the sgNT DMSO (N=1) and rapamycin-treated (N=2) groups. Taken together, these findings suggest that a rapamycin-based induced proximity approach could be used to identify genes that BAF can activate.

Following the execution of the screen, we found that sgRNAs spanning 79 genes were significantly depleted (P <0.05, Log_2_[Fold Change] <-0.6), suggesting that the potential activation of these genes by BAF has proliferation suppressive properties in the context of K562 cells (**Figure 4B**). Interestingly, only 5% of these hits are previously known tumor suppressors^57^ (**Figure 4B**). To further understand the biological function of these BAF-activated genes, Gene Set Enrichment Analysis^58,59^ (GSEA) was performed on all genes ranked by MAGeCK analysis^60^. Interestingly, genes that inhibit cellular proliferation when activated by BAF are involved in the hypoxia pathway and the TGFβ signaling pathway (**Figure 4C, Supplemental Table 1**). On the contrary, genes that induce cellular proliferation when activated by BAF are involved in the bile acid metabolism pathway and the TNFα signaling via NFκB pathways (**Figure 4C, Supplemental Table 1**).

To understand the epigenetic signatures that predict responsiveness to BAF-mediated gene activation, we compared the top 500 BAF-mediated activated genes with publicly available ChIP seq datasets for the active H3K4me3 and H3K27Ac epigenetic marks, and the repressive H3K27me3 epigenetic marks (GSE108323^61^ and GSE128262^62^). Interestingly, about 46.6% (223/500) of genes activated by BAF have dual active H3K4me3 and H3K27Ac marks at their promoters (**Figure 4D**). 17.6% (88/500) of genes activated by BAF have neither of these active or repressive epigenetic marks, while 13% of genes activated by BAF contained both active epigenetic marks (H3K4me3, and H3K27Ac) and the repressive H3K27me3 epigenetic mark. In addition, 13.2% (66/500) of genes only contain the activating H3K27Ac mark (**Figure 4D**). Only 3.2% of genes activated by BAF are bivalent (**Figure 4D**). The combination of the active H3K4me3 and repressive H3K27me3 marks define bivalent promoters that are poised to activate transcription upon exposure to appropriate stimuli. Taken together, this suggests that the BAF complex has the largest propensity to induce expression in genes with dual activation marks on their promoters. In contrast, our data suggests BAF has limited capacity to activate bivalent genes with H3K4me3 and H3K27me3 makes on their promoters.

To further understand the ability of BAF to activate genes, we compared the top 500 activated genes with their basal RNA expression levels and ranked genes into ten deciles by their baseline expression. We discovered that BAF-mediated activation does not discriminate based on a gene’s basal expression level, activating genes with high and low basal expression levels (**Figure 4E**). Overall, BAF appears to activate genes with different basal levels of expression and epigenetic states, highlighting BAF’s robustness in driving gene expression.

### BAF activates novel genes that have the potential to suppress proliferation

As mentioned previously, in addition to known tumor suppressors, we found that BAF could activate novel genes that inhibited cell proliferation (**Figure 4B**). Among the significantly depleted genes, we focused on *SDF2* and *SPI1* as potential candidates to validate our screen and the robustness of the BAF complex as a transcriptional activator (**Supplemental Figure 4A**). *SDF2,* or stromal-derived factor 2, is an ER-resident protein where high transcript levels are associated with good prognosis in breast cancer, while low levels of SDF2 are associated with shorter disease-free survival in colorectal carcinomas^63,64^. *SPI1* or *PU.1* is an ETS-domain transcription factor essential in driving myeloid and B-lymphoid development^65,66^. Depending on the disease context, *SPI1* can function as an oncogene^67,68^ or a tumor suppressor^69,70^.

To validate the effects of *SDF2* and *SPI1* on cell viability, we transduced K562-FIRE cells with specific sgRNAs targeting the promoters of these genes (**Figure 5A-B**). Next, we monitored cellular proliferation following rapamycin treatment to induce BAF recruitment. Cell growth inhibition was initially observed in cells transduced with both targeting sgRNA and non-targeting sgRNA (sgNT) (**Figure 5C-D**), but this decrease in cell proliferation can be attributed to the pharmacological effects of rapamycin^71^. After day 7, the tumor suppressive effect of activating these genes becomes more apparent. By day 14, BAF-mediated activation of *SDF2* resulted in a significant decrease in cell growth when compared to sgNT (**Figure 5C**). Similarly, rapamycin-treated cells expressing with *SPI1* sgRNAs also showed decreased cell proliferations, especially in cells expressing sg*SPI1* 3 (**Figure 5D**).

**Figure 5.**
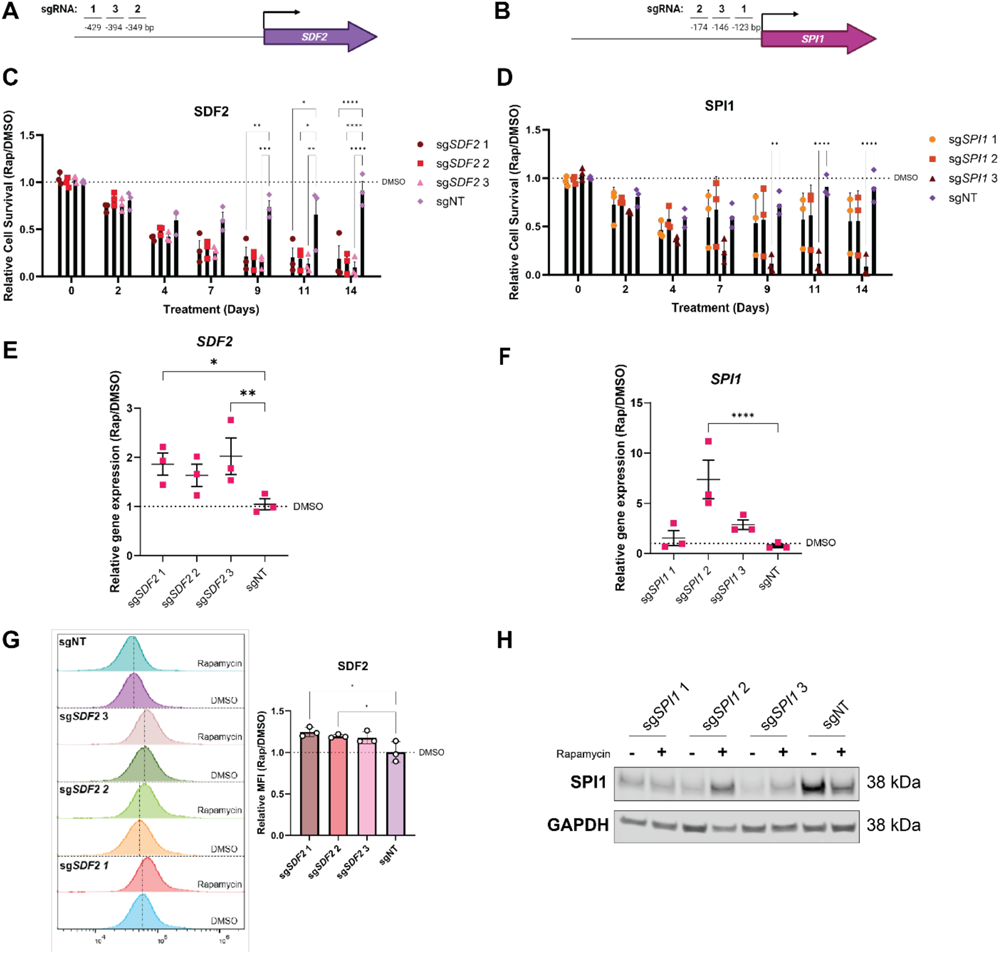
BAF-mediated activation of cell proliferation suppressors highlights the potential for targeted therapeutics. **(A)** Schema of the *SDF2* promoter with corresponding sgRNA binding locations upstream of the transcriptional start site. **(B)** Schema of the *SPI1* promoter with corresponding sgRNA binding locations upstream of the transcriptional start site. **(C)** Relative cell survival of K562-FIRE cells when *SDF2* is activated by BAF after 10 nM of rapamycin treatment. **(D)** Relative cell survival of K562-FIRE cells when *SPI1* is activated by BAF after 10 nM of rapamycin treatment. **(E)** Expression of *SDF2* in K562-FIRE cells expressing sg*SDF2* or sgNT after 48 h of 10 nM rapamycin or DMSO treatment. **(F)** Expression of *SPI1* in K562-FIRE cells expressing sg*SPI1* or sgNT after 48 h of 10 nM rapamycin or DMSO treatment. **(G)** Representative flow cytometric profile and MFI of SDF2 in K562-FIRE cells expressing sg*SDF2* or sgNT after 48 h of 10 nM rapamycin or DMSO treatment. **(H)** Representative immunoblot of SPI1 in K562-FIRE expressing sg*SPI1* or sgNT were after 48h of 10 nM rapamycin or DMSO treatment. All experiments were shown from 3 independent experiments with qPCR and cell survival performed in triplicates. Data are presented as mean ± SEM. Significance was determined by 1-way ANOVA with Turkey’s multiple-comparisons test, comparing targeting sgRNA with sgNT. \**P* < 0.05, \*\**P* < 0.01, \*\*\**P* < 0.001, \*\*\*\**P* < 0.0001

Furthermore, in comparison to sgNT, recruiting the BAF complex to these promoters increased *SDF2* (~2-fold increase in sg*SDF2* 1 and 3) and *SPI1* (~8-fold and 3-fold increase in sg*SPI1* 2 and 3 respectively) mRNA levels (**Figure 5E-F**). Concomitantly, BAF-mediated activation of *SDF2* led to a subtle increase of SDF2 protein when assessed using intracellular FACS analysis (**Figure 5G**). This is intriguing as a small change observed in the mRNA and protein levels of SDF2 has a profound effect on cell viability. This suggests a possible role of SDF2 in maintaining a tight balance between cell survival and cell death cellular programs, such that offsetting this balance would result in cell death^72,73^. In addition, BAF-mediated activation of *SPI1* also led to an increase in SPI1 protein levels, reflecting the observed increase in *SPI1* mRNA levels (**Figure 5F**). Collectively, this data validates our screen, confirming its ability to identify genes that, when activated by BAF’s recruitment to the gene’s promoter, results in the inhibition of cellular proliferation.

Finally, this work highlighted the transcriptional potential of the BAF complex via induced proximity. The BAF complex can be recruited to promoters, increase chromatin accessibility, and change local epigenetic status to drive gene expression. Our work also shows the viability of using such induced proximity modality to harness BAF complexes to activate genes with inherent anti-proliferative effects, laying the foundation for developing future induced proximity drugs for gene activation.

## Discussion

Transcription regulation is a complex process involving regulatory elements, transcription factors, chromatin dynamics, and epigenetic changes. The BAF complex promotes gene expression by using energy from ATP hydrolysis to remodel chromatin, providing access to transcriptional machinery^11^. Overexpression of BAF subunits in BAF null cancer cells increases chromatin accessibility at enhancers, driving the expression of active and bivalent genes^21–23^. Using CIP, targeted recruitment of BAF via zinc-finger proteins or dCas9 results in increased local chromatin accessibility and a corresponding increase in gene expression, particularly in select bivalent genes in mouse embryonic stem cells^25,26^. In our study, we leveraged the highly adaptable FIRE-dCas9 system in human cellular models and tested its ability to activate therapeutically relevant genes in K562 and 293T cells. Our ChIP analysis validated successful BAF recruitment to the *HBG1/2* promoter and delineated the engagement of cBAF and ncBAF to promoter regions (**Figure 1**). This engagement increased chromatin accessibility, consequently driving *HBG1* expression (**Figure 1–2**). Contrary to previously established models, targeted BAF recruitment to the enhancers of *HBG* did not significantly increase the expression of globin genes (**Supplemental Figure 2**). This is interesting as pharmacological inhibition of the BAF complex in various cellular models abrogates BAF binding and function at the enhancers^38,40,74^. Natively, the various BAF complexes have distinct genomic distribution patterns, reflecting their specific functions^75^. cBAF and PBAF predominantly localize to the enhancers and promoters, respectively, while ncBAF localizes to promoters and genomic regions bound by CTCF, suggesting its additional role in regulating transcription through chromosomal looping^5^. While our study demonstrated that redirecting cBAF and ncBAF to the promoters activates transcription, the underlying mechanism remains unclear. To elucidate this, the FIRE-dCas9 system could be adapted to recruit individual distinct BAF complexes or collectively to target the enhancers and promoters. This approach would enhance our understanding of BAF’s specific role in transcriptional activation.

In highlighting the promoter-centric transcriptional activity of the BAF complex, our study uncovered BAF’s nuanced role in modulating local epigenetic states when directed to the *HBG1/2* promoter. Previous studies demonstrated that targeted BAF recruitment to an engineered *Pou5f1* locus in mouse embryonic fibroblast increases chromatin accessibility while evicting PRC1 and PRC2 complexes to establish a permissive environment for transcription^26,51^. Similarly, when BAF was targeted to the promoter of the neuro-specific *Nkx2.9* bivalent gene in mouse embryonic stem cells, a substantial increase in its expression was observed, which was accompanied by a subtle decrease in repressive H3K27me3 marks and an enrichment of activating H3K4me3 marks at the promoter^25^. In our study, we found that targeting BAF to the *HBG* loci enriches H3K4me3 marks in K562 cells and H3K27Ac marks in 293T cells (**Figure 3A-B**). Furthermore, targeted BAF recruitment significantly depleted H2AK119Ub (PRC1) and H3K27me3 (PRC2) marks in 293T cells (**Figure 3C**). Collectively, data from previous studies and our work here highlight some commonalities in epigenetic changes observed upon BAF recruitment.

The ability of the FIRE-dCas9 system to target specific genomic loci adds to the current CRISPR/Cas9 toolbox^76^, allowing one to probe the biology of any protein complex that has the potential to modulate chromatin dynamics and epigenetic states. Using this approach, previous studies have shown that targeted recruitment of BAF to bivalent genes induced gene expression^25^. Consistent with this, the overexpression of BAF in cell lines lacking BAF results in the activation of multiple bivalent genes^21–23^. These observations suggest determining a promoter’s bivalency could predict its responsiveness to BAF-mediated activation. However, a comparative analysis of BAF-activated genes identified through our genome-wide CRISPR screen preliminarily suggests that factors beyond bivalency may govern the responsiveness to BAF-mediated activation. When considering native epigenetic marks and basal expression levels, our findings indicate that BAF primarily activates genes containing H3K4me3 and H3K27Ac marks on their promoters, and interestingly, its ability to activate genes does not seem to be dependent on their basal expression level (**Figure 4D-E**). Thus, to identify predictive factors governing responsiveness to BAF-mediated activation, additional studies such as single-cell RNA-sequencing, ATAC-sequencing, and ChIP-sequencing using the FIRE-dCas9 system could prove useful. These efforts will provide an in-depth understanding of the biology of targeted BAF-mediated activation, allowing us to predict genes that BAF can activate.

Induced proximity is integral to various biological processes, including receptor function, transcriptional and translational functions, protein localization, and stabilization^28^. More recently, efforts have been focused on developing heterobifunctional molecules to induce proximity chemically^27,77^. A successful example of this is the recruitment of a target protein of interest to the proteasome for degradation or inhibiting protein function by stabilizing protein-protein interactions^27^. More recently, transcription chemical inducers of proximity (TCIP), potent heterobifunctional molecules, have been developed as a novel approach to induce cell death^78,79^. The TCIP molecules achieve this by redirecting endogenous transcription activators such as BRD4 or CDK9 to transcriptionally repressed BCL6 sites, thereby activating apoptotic pathways in diffuse large B-cell lymphoma (DLBCL)^78–80^. The recruitment of these transcriptional activators is sufficient to overcome the transcriptional repressive effects of BCL6, establishing a gain-of-function mechanism to activate apoptotic genes resulting in cell death, suggests that a small molecule heterobifunctional transcriptional activator, utilizing induced proximity, could be developed as a therapy^78,79^. Our study builds on this, demonstrating that BAF can act as a transcriptional activator to increase the expression of genes with therapeutic potential via CIP (**Figure 5**). Due to the modularity of these heterobifunctional molecules, BAF could potentially be recruited via CIP to proteins residing at the loci of interest^78,79^. To identify proteins bound to a gene of interest, the FIRE-dCas9 system could be modified to incorporate a dCas9 protein fused to a biotin ligase^81,82^. This modification would enable the labeling and identification of proteins in the vicinity via mass spectrometry. One can then develop binders to these proteins and, ultimately, heterobifunctional molecules with the potential to recruit BAF to these specific loci of interest.

Our study additionally highlights a novel therapeutic strategy for treating beta-globinopathies by elevating fetal hemoglobin levels. Despite extensive research into small molecule therapies inducing fetal hemoglobin expression, clinical evidence of their safety and efficacy remains limited^83^. Current clinical approaches involve hydroxyurea, short-chain fatty acids like 2,2-dimethylbutyrate, and DNA methylation inhibitors such as 5-azacytidine or Decitabine^83^. Recent advances in gene editing CRISPR/Cas9 technologies have been leveraged to treat sickle cell disease genetically, either by correcting the disease-causing mutation in adult hemoglobin or by increasing fetal hemoglobin levels through editing regulatory elements or inhibiting known repressors such as BCL11A^84,85^. As a potential additional therapeutic approach for beta-globinopathies, our study reveals that when BAF is recruited to the *HBG1/2* promoter, it induces fetal hemoglobin expression at both the mRNA and protein levels (**Figure 2**). Furthermore, given that repressor proteins like BCL11A and KLF1 have been shown to bind to the *HBG* promoter to suppress its expression^41,53,84^, one can consider the development of a heterobifunctional molecule that recruits BAF to the *HBG* loci via these repressor proteins to induce fetal hemoglobin expression, offering a novel approach for treating beta-globinopathies. In conclusion, our study lays the foundation for developing a BAF-centric heterobifunctional molecule to activate therapeutically relevant genes via induced proximity.

## Materials and Methods

Detailed materials and methods are provided in the supplemental material, including culture medium, sgRNA, primers, and antibodies (**Supplemental Table 2-4**). RNA-sequencing and CRISPR screening data were deposited into the NCBI Gene Expression Omnibus (GSE249747). Complete unedited blots are found in the supplemental material.

## Acknowledgments

We want to thank our many Foghorn Therapeutics colleagues for their unwavering support and help through the project, especially Qianhe Zhou, Sal Topal, Julie Di Bernardo, Dave Lahr, and David Mayhew for providing valuable insights for this work.

## Author contribution

L.C. and A.T. conceptualized and designed the study. L.C. and B.S. performed the experiments and interpreted the data. D.M. and D.L. performed bioinformatics analysis and interpretation of the RNA-seq and CRISPR screen results. L.C. wrote the manuscript, with D.M., B.S., D.L., and A.T. providing revisions before submission.

