## Supplemental Figure 1 for "Hijacking the transcriptional activation potential of the BAF complex via Induced Proximity"

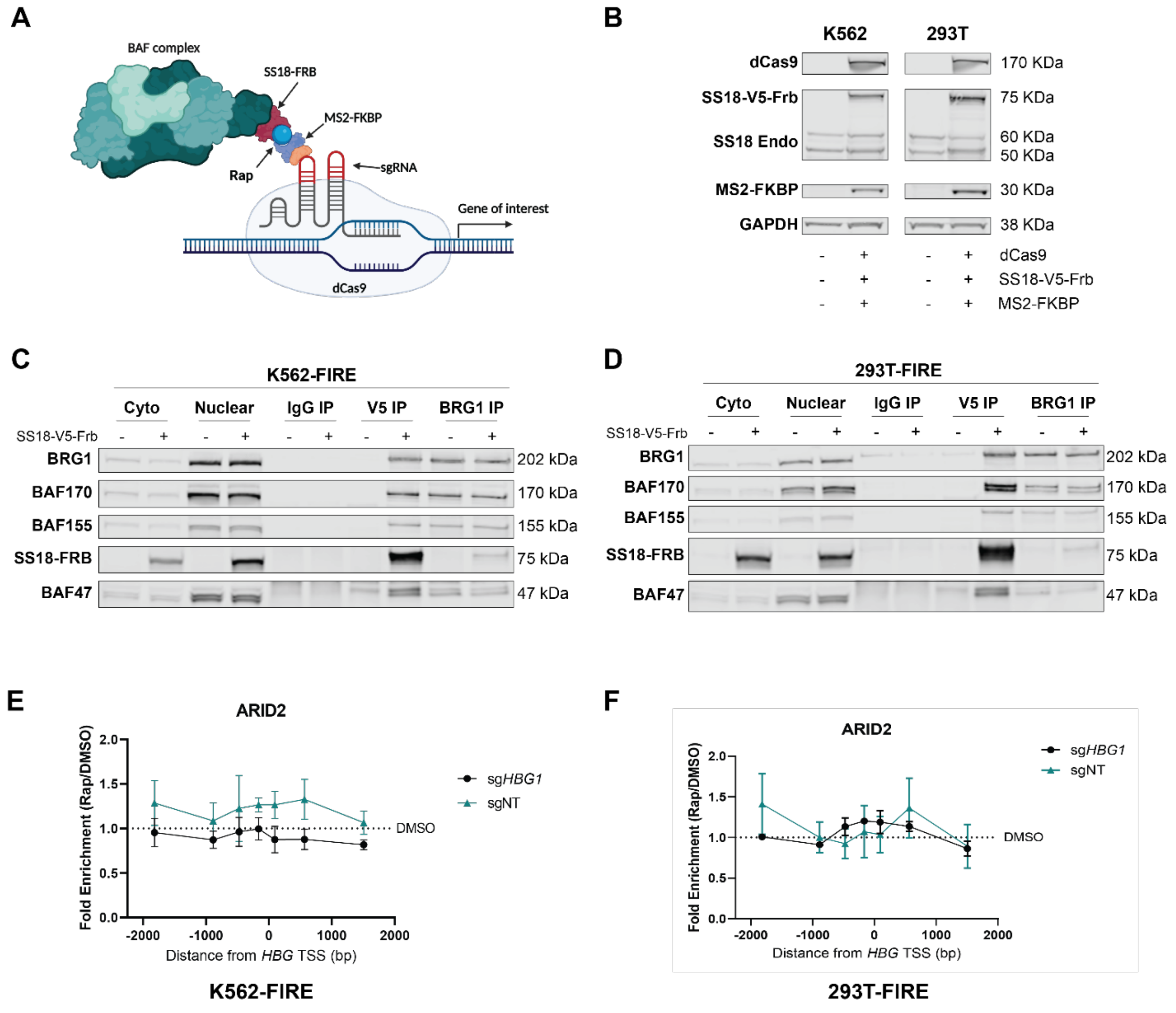


**Supplemental Figure 1. Rapamycin inducible SS18-V5-FRB system is ectopically expressed and incorporated into the BAF complex.**

- **(A)** A schematic representation of the FKBP/FRB inducible recruitment for epigenome editing by Cas9 (FIRE-dCas9) system. Created with BioRender.
- **(B)** Representative immunoblot of the FIRE-dCas9 system in K562 and 293T cells.
- **(C)** Representative co-immunoprecipitation (Co-IP) immunoblot of SS18-V5-FRB interacting with BAF subunits in K562-FIRE cells.

**(D)** Representative Co-IP immunoblot of SS18-V5-FRB interacting with BAF subunits in 293T-FIRE cells.

**(E)** ChIP-qPCR comparing PBAF complex occupancy (ARID2) across the *HBG1* loci in K562-FIRE cells expressing sg*HBG1* or sgNT after 24 h of 10 nM rapamycin or DMSO treatment.

**(F)** ChIP-qPCR comparing PBAF complex occupancy (ARID2) across the *HBG1* loci n 293T-FIRE cells expressing either sg*HBG1* or sgNT after 24 h of 10 nM or DMSO treatment.

Data shown are representative of 3 independent experiments. All data shown for ChIP-qPCR experiments are from 3 independent experiments performed in triplicate. Data are presented as mean ± SEM. Significance was determined by 1-way ANOVA with Turkey’s multiple-comparisons test, with indicated significance showing the comparisons between sg*HBG1* and sgNT.
