## Supplemental Figure 2 for "Hijacking the transcriptional activation potential of the BAF complex via Induced Proximity"

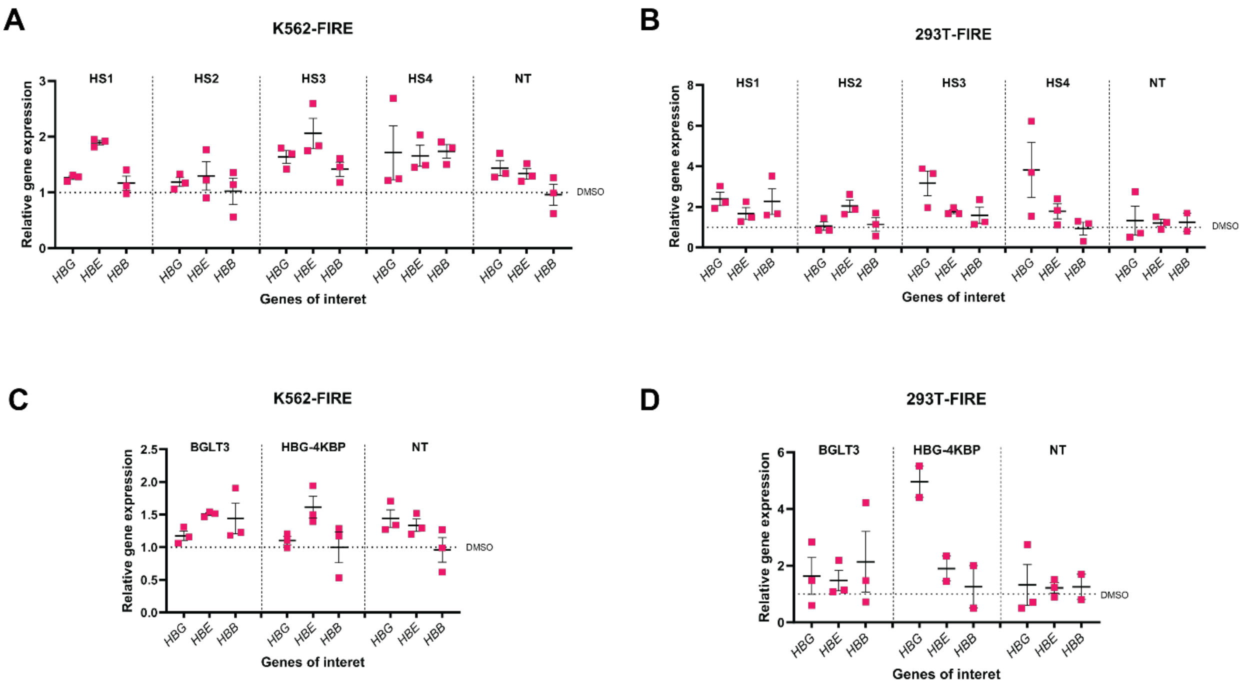


**Supplemental Figure 2. Targeted recruitment of BAF to the enhancers of HBG1/2 does not induce mRNA expression.**

**(A)** Expression of globin genes *HBG1,* embryonic hemoglobin (*HBE*), and adult hemoglobin (*HBB*) when BAF is targeted to enhancer regions HS1-4 using sgRNAs. Expression was determined 48 h after 10 nM rapamycin or DMSO treatment in K562-FIRE cells.

**(B)** Expression of globin genes *HBG1, HBE,* and *HBB* when BAF is targeted to enhancer regions HS1-4 using sgRNAs. Expression was determined 48 h after 10 nM rapamycin or DMSO treatment in 293T-FIRE cells.

**(C)** Expression of globin genes *HBG1, HBE*, and *HBB* when BAF is targeted to novel enhancer regions at the *BGLT3* gene locus and the -4 kbp region upstream of the *HBG1* promoter. Expression was determined 48 h after 10 nM rapamycin treatment in K562-FIRE cells.

**(D)** Expression of globin genes *HBG1, HBE,* and *HBB* when BAF is targeted to novel enhancer regions at the *BGLT3* gene locus and the -4 kbp region upstream of the *HBG1* promoter.

Expression was determined 48 h after 10 nM rapamycin treatment in K562-FIRE cells. Data shown from at least two independent experiments performed in triplicate.

Data shown from 3 independent experiments performed in triplicates. Data are presented as mean ± SEM. Significance was determined by 1-way ANOVA with Turkey's multiple-comparisons test, comparing sg*HBG1* with sgNT.
