## Supplemental Figure 3 for "Hijacking the transcriptional activation potential of the BAF complex via Induced Proximity"

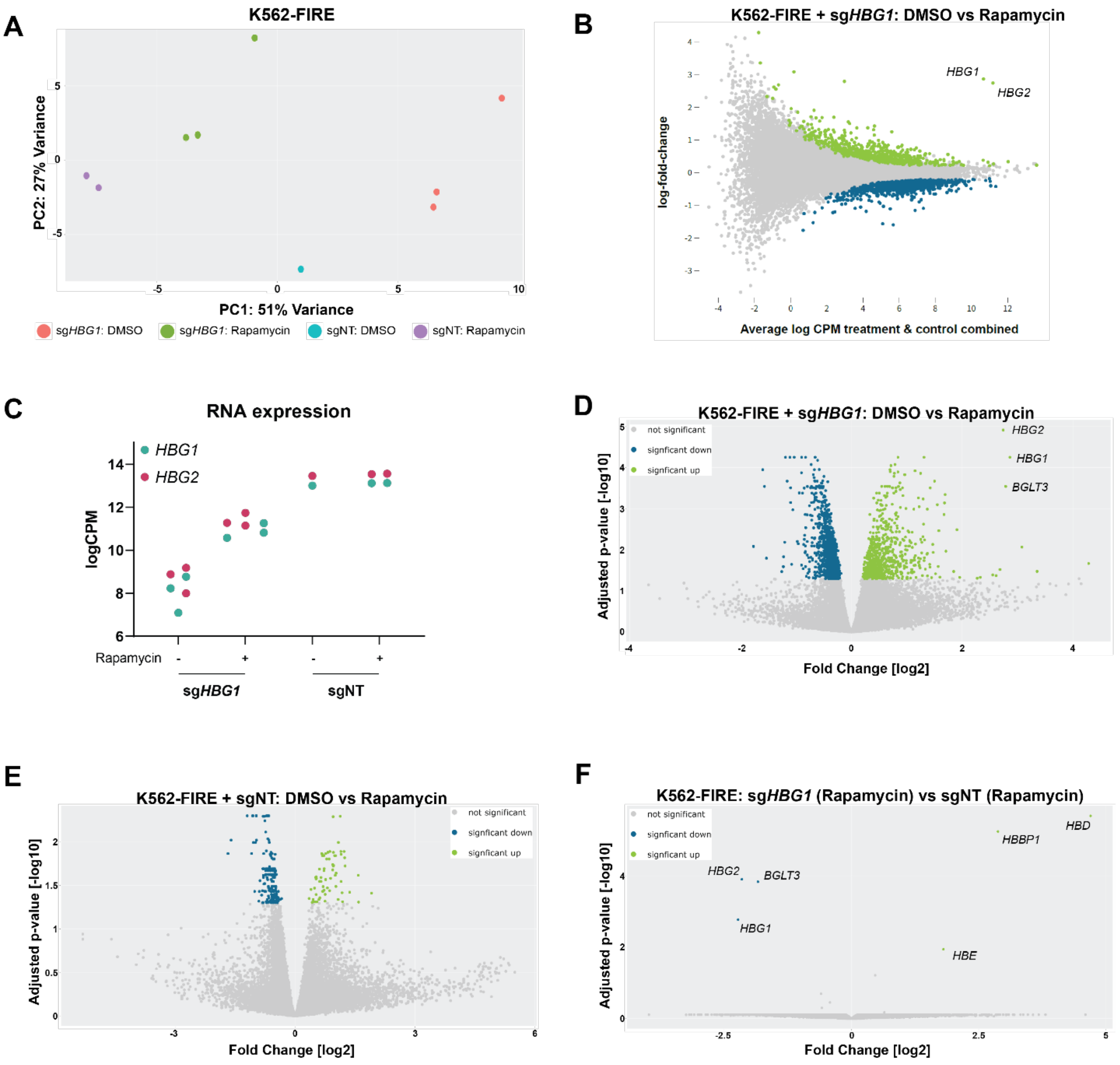


**Supplemental Figure 3. Expression of *HBG1/2* is specific when utilizing the FIRE-dCas9 system.**

**(A)** Principal component analysis of RNA-seq performed in K562-FIRE cells expressing sg*HBG1* or sgNT after 24 h of 10 nM of rapamycin or DMSO treatment. Due to low-quality RNA extractions, limited sgNT samples were sequenced (sgNT – DMSO: N=1; sgNT – Rapamycin: N=2)

**(B)** Mean difference plots of K562-FIRE cells expressing sg*HBG1* treated after 24 h of 10 nM rapamycin or DMSO.

**(C)** *HBG1/2* transcript levels in K562-FIRE cells expression sg*HBG1* or sgNT after 24 h of 10 nM rapamycin or DMSO treatment.

**(D)** Volcano plot of differentially expressed genes (DEG) in K562-FIRE cells expressing sg*HBG1* after 24 h of 10 nM rapamycin or DMSO treatment. 1,075 genes were significantly downregulated, while 690 genes were significantly upregulated when rapamycin-treated cells were compared to DMSO.

**(E)** Volcano plot of DEGs in K562-FIRE cells expressing sgNT after 24 h of 10 nM rapamycin or DMSO treatment. 175 genes were significantly downregulated while 67 genes were significantly upregulated when rapamycin-treated cells were compared to DMSO-treated cells.

**(F)** Volcano plot of DEGs in rapamycin-treated K562-FIRE cells expressing sg*HBG1* or sgNT. Only three genes were upregulated and downregulated when rapamycin-treated cells expressing sg*HBG1* were compared to rapamycin-treated cells expressing sgNT after 24h of treatment.

Data shown are from 3 independent replicates in sg*HBG1* (Rapamycin and DMSO), 2 independent replicates in sgNT (rapamycin), and 1 experiment in sgNT (DMSO) group.
