## Supplemental Figure 4 for "Hijacking the transcriptional activation potential of the BAF complex via Induced Proximity"

**A**

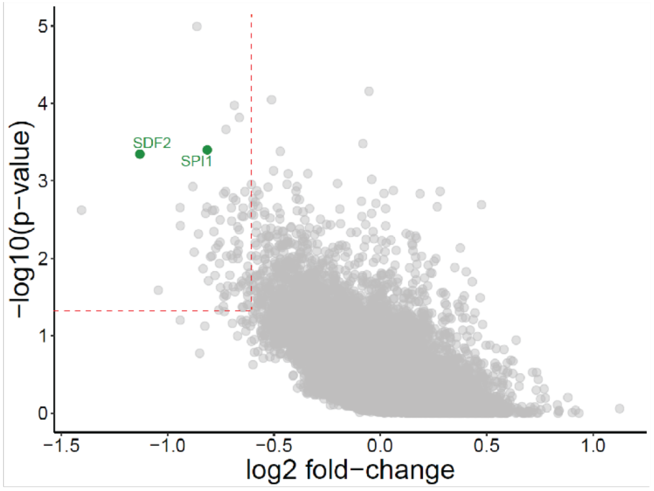


| Gene | P-value | LogFC | H3K27Ac | H3K27me3 | H3K4me3 | Expression Decile |
|---|---|---|---|---|---|---|
| *SDF2* | 0.000452 | -1.1304 | ✓ | × | ✓ | 9th |
| *SPI1* | 0.0004 | -0.81384 | ✓ | × | ✓ | 8th |

**Supplemental Figure 4. Selected genes that exhibit a highly significant loss in sgRNA in whole-genome BAF-mediated activation screen.**

**(A)** P-values and log Fold Changes of selected genes that BAF activates. Also shown are the corresponding baseline epigenetic marks on their promoters and their baseline expression level (Decile 1 being the lowest expressing and decile 10 being the highest expressing genes). Dashed lines indicate thresholds of $P < 0.05$ and $Log_2$[Fold Change] < -0.6.
