## Supplemental Information for "Hijacking the transcriptional activation potential of the BAF complex via Induced Proximity"

### Hijacking the transcriptional activation potential of the BAF complex for therapeutic purposes via Induced Proximity

#### Supplemental information

1. **Materials and Methods**
2. **Supplemental Table 2.** sgRNA sequences determined by the Broad sgRNA database and konermann *et al.*
3. **Supplemental Table 3.** Primers used for ChIP-qPCR and ATAC-qPCR
4. **Supplemental Table 4.** Antibodies used for immunoblotting, Co-IP, ChIP, and Flow cytometry
5. **Uncut gels**

#### Material and Methods

##### Cell lines

K562, a chronic myelogenous leukemia cell line (American Type Culture Collection, ATCC; CCL-243) was cultured in Iscove's Modified Dulbecco's Medium (Gibco, ThermoFisher Scientific) supplemented with 1X GlutaMAX (Gibco, ThermoFisher Scientific) and 10% FBS (Omega Scientific). HEK293T, a human embryonic kidney cell line (ATCC), was cultured in Dulbecco's Modified Eagle Medium supplemented (Gibco, ThermoFisher Scientific) with 1X GlutaMAX (Gibco, ThermoFisher Scientific) and 10% FBS (Omega Scientific). Cells were routinely tested for mycoplasma and validated for STR.

##### Plasmids and CRISPR sgRNA plasmids

Plasmids encoding the FIRE-dCas9 system[1] (Lv EF1a dCas9 2A Blast; EF1a MS2-Fkbp(1x) 2A Hygro; Lv EF1a SS18-Frb PGK Puro) were synthesized and cloned into pGen_Lenti (GenScript). sgRNA used to target our genes of interest were designed using the Broad sgRNA Database (GenScript) and cloned into SAM gRNA plasmids (GenScript) (**Supplemental Table 2**). Additional sgRNA sequences targeting enhancer regions were designed using CRISPick[2,3].

##### Lentivirus production and delivery

To generate cell lines stably expressing the FIRE-dCas9 system, lentiviral plasmids, sand packaging plasmid mix (Cellecta) were co-transfected into HEK293T cells using Lipofectamine 2000 (ThermoFisher Scientific) according to manufacturer's directions. 48 h post-transfection, viral supernatant was centrifuged at 2,500 x g for 10 min at 4 °C to remove cell debris. Viral supernatant was mixed with PEG-it concentration reagent (System Biosciences) and incubated overnight at 4 °C. The mixture was centrifuged at 1,500 x g for 30 mins at 4 °C. The Virus pellet was resuspended in PBS, and the virus titer was measured using QuickTiter Lentivirus Titer Kit (Cell Biolabs, Inc.) according to manufacturer's protocol. K562 cells and 293T cells were transduced with these viruses at a Multiplicity of infection (MOI) of 0.5 in the presence of polybrene (8 µg/mL, Millipore Sigma) and selection of these constructs was achieved with blasticidin (K562: 10 µg/mL; 293T: 12 µg/mL), hygromycin (K562: 200 µg/mL; 293T: 200 µg/mL) and puromycin (K562: 3 µg/mL; 293T: 1 µg/mL). CRISPR sgRNA lentivirus was generated similarly, and transformed cells were selected with zeocin (K562: 200 µg/mL; 293T: 400 µg/mL).

**RNA isolation, reverse transcription, and quantitative PCR analysis (qPCR)**

After cells were treated with DMSO or rapamycin, total RNA was isolated from cells using RNAeasy Mini Kit (Qiagen) according to manufacturer's protocol. cDNA was synthesized from isolated RNA using the High-capacity cDNA reverse transcription kit (Applied Biosystems). Quantitative PCR was used to assess changes in expression using Taqman probes (ThermoFisher Scientific): *HBG1/2* (Hs00361131_g1), *HBE* (Hs00362215_g1), *HBB* (Hs00758889_s1), *SDF2* (Hs01054851_g1), *SPI1* (Hs02786711_m1), and *GAPDH* (Hs99999905_m1). qPCR was performed on the QuantStudio 7 Flex (ThermoFisher Scientific), and relative expression was determined by $2^{-\Delta\Delta CT}$ with *GAPDH* as a loading control.

**Assay for transposase-accessible with qPCR (ATAC-qPCR)**

DNA isolated from cells for ATAC-qPCR was performed using the ATAC-Seq Kit (Active Motif according to manufacturer's protocol. In brief, isolated cells were washed with cold PBS, lysed with ice-cold ATAC lysis buffer (Active Motif), and centrifuged at 500 x g for 10 mins 4 °C. Supernatant is carefully removed, and Tagmentation mix (1X Tagmentation buffer, 0.4X PBS,

0.01% Digitonin, 0.1% Tween 20, and assembled transposons) is added to the nuclear pellets. The reaction is incubated at 37 °C for 30 mins in a thermomixer at 800 rpm, and tagmented DNA is purified using the MinElute PCR purification kit (Qiagen) according to manufacturer's protocol. The tagmented DNA is PCR amplified using provided indexing primers, and the resulting DNA is purified using the MinElute PCR purification Kit (Qiagen). Accessibility of chromatin along the *HBG1* loci was assessed by qPCR using PowerUp SYBR Green master mix (ThermoFisher) performed on CFX384 (Bio-Rad). Enrichment at specific genomic regions is compared to a control region without BAF binding, lack of active epigenetic marks, and low ATAC accessibility using the $2^{-\Delta\Delta CT}$ method (Primers listed in **Supplemental Table 3**).

**Chromatin immunoprecipitation with qPCR (ChIP-qPCR)**

ChIP was performed as described previously[4]. In brief, isolated cells ($4x10^6$ cells / ChIP) were washed with PBS and fixed for 10 mins using 1% formaldehyde (Sigma-Aldridge) at room temperature for 10 min while shaking. Glycine solution (1X, Cell Signaling Technologies) was added to quench the crosslinking reaction, and cells were incubated for 10 mins at room temperature while shaking. Cells were subsequently washed twice in cold PBS, and nuclear pellet was isolated by lysing cells with cell lysis buffer (20 mM Tris pH 8.0, 85 mM KCl, and 0.5% NP40) containing Halt Protease inhibitors (ThermoFisher Scientific) and centrifuged at 450 x g for 5 mins at 4 °C. Nuclei pellets were collected, lysed with nuclei lysis buffer (10 mM Tris-HCl pH 7.5, 0.5 % Sodium Deoxycholate, 0.1 % SDS, and 1 % NP40) containing Halt Protease inhibitors (ThermoFisher Scientific). Chromatin is sheared in a water bath sonicator for 20 mins at 40 % amplitude with 15 sec on and 15 sec off (Qsonica Sonicators). Sheared chromatin was spun at 18,000 x g for 10 mins at 4 °C. Supernatant was collected, and an aliquot of sheared chromatin was used as input.

Sheared chromatin was immunoprecipitated, incubating chromatin with antibodies and 30 μL of Protein A/G Dynabeads (1:1 mix, ThermoFisher Scientific) overnight at 4 °C while rotating (**Supplemental Table 4**). Immunoprecipitates were washed four times with wash buffer (10 mM Tris-HCl pH 7.5, 1 % NP40, and 0.1 % SDS) and eluted with 100 μL of elution buffer (50 mM sodium bicarbonate and 1 % SDS). Immunoprecipitated DNA was reverse crosslinked with 50 mM NaCl, incubated overnight at 65 °C, and treated with RNase A (ThermoFisher Scientific) and Proteinase K (ThermoFisher Scientific). Immunoprecipitated DNA was purified using the MinElute PCR purification kit (Qiagen) according to manufacturer's protocol, and purified ChIP DNA was analyzed by performing qPCR using PowerUp SYBR Green master mix (Applied Biosystems) on the CFX384 (Bio-Rad) (Primers are listed in **Supplemental Table 3**). Changes in enrichment are calculated by normalizing enrichment (bound/input) in rapamycin-treated groups to those in the DMSO-treated group (Rap/DMSO).

**RNA sequencing (RNAseq)**

After $1 \times 10^6$ cells were treated with DMSO or rapamycin, cells were pelleted, and RNA was isolated and sequenced (Novogene) with paired-end 150 bp reads to a depth of ~20 million reads per sample. Reads were aligned to the human genome (GRCh38) using STAR v2.7.3a, and gene counts were quantified on the GENCODE reference gtf file version 27 using featureCounts from the SubRead package (v2.0.0). Differential gene expression was calculated from the resultant gene-level counts matrix using the limma-voom method (after TMM normalization) in R (R v4.1.2, limma v3.34.9, Glimma 1.10.0, and edgeR v3.24.0). Statistically significant genes were those with an FDR-corrected p-value less than 0.05. Gene set enrichment analysis (GSEA) was performed with the MSigDB[5,6] (h.all.v2023.1.Hs.symbols.gmt) gene sets using the R package clusterProfiler (v4.2.2).

**CRISPRa sgRNA library lentivirus production and transductions**

The SAM CRISPRa sgRNA library was obtained from GenScript. CRISPRa library lentivirus was generated according to manufacturer's protocol (Cellecta). In brief, HEK293T cells were transfected with the CRISPRa library and the packaging plasmid mix (Cellecta) using Lipofectamine 2000 (ThermoFisher Scientific). 48 h post-transfection, viral supernatant was centrifuged at 2,500 x g for 10 min at 4 °C to remove cell debris. Viral supernatant was mixed with PEG-it concentration reagent (System Biosciences) and incubated overnight at 4 °C. The mixture was then centrifuged at 1,500 x g for 30 mins at 4 °C, and the virus pellet was resuspended in PBS. Functional viral titers in K562 cells were determined using the Global UltraRapid Lentiviral Titering Kit (System Bioscience) according to manufacturer's protocol. K562-FIRE cells were then transduced with the library at a low MOI of 0.3 while still ensuring coverage of >500 sgRNA per gene per guide. Cells were spinfected with polybrene (8 µg/mL, Millipore Sigma) at 2,000 x rpm for 2 h at room temperature, following which spinfected cells were incubated for 24 h in a 37 °C incubator. Cells were selected for the library with zeocin (200 µg/mL) for at least 14 days.

**CRISPR screen analysis**

After establishing the library expressing K562-FIRE cells, 1 x $10^8$ cells were treated with either DMSO or rapamycin for 14 days, with refreshment of rapamycin and media after 2-3 days. 1 x $10^8$ cells were collected from each treatment for genomic DNA isolation and deep sequencing (Cellecta). This screen is performed in triplicate from 3 independent transductions. Trimmed fastq reads were aligned to an index of the sgRNA library sequences using bowtie (v1.3.1) with no mismatches allowed. sgRNA counts were analyzed with MAGeCK (v0.5.9.2)[7] using the 'Mageck test' RRA to rank the genes under negative selection ins the rapamycin treatment over DMSO. MAGeCK's negative selection gene ranking was used for Gene set enrichment analysis (GSEA), which was performed with the MSigDB[5,6] (h.all.v2023.1.Hs.symbols.gmt) gene sets using the R package clusterProfiler (v4.2.2).

**Immunoblotting**

Total proteins were isolated from cells using RIPA buffer (Millipore Sigma) containing Halt protease and phosphatase inhibitor cocktail (ThermoFisher Scientific). 30 μg of lysates were separated on a gradient (4-12%) bis-tris gel (ThermoFisher Scientific) and transferred onto a nitrocellulose membrane (ThermoFisher Scientific) through a semi-dry transfer using the P0 program on the iBlot 2 (ThermoFisher Scientific). Blots were probed with various primary antibodies (**Supplemental Table 4)**, and protein bands were visualized using the Odyssey CLx imager (LI-COR Biosciences). GAPDH was used as a loading control.

**Immunoprecipitation and Co-immunoprecipitation**

Isolated protein (0.75 mg) was used to enrich HBG in K562-FIRE cells after treatment with DMSO and rapamycin for 72 h. Lysates were incubated overnight at 4 °C with 30 μL of Protein A/G beads (ThermoFisher Scientific) coated with 3 μg of HBG or IgG antibody (**Supplemental Table 4**). Beads were washed, and immunoprecipitates were separated on a gradient bis-tris gel (4-12%), transferred onto a nitrocellulose membrane (ThermoFisher Scientific) and probed with HBG and GAPDH antibodies. Protein bands were visualized on the Odyssey CLx imager (LI-COR Biosciences) (**Supplemental Table 4**). Similarly, to co-immunoprecipitate protein interacting protein partners, nuclear lysates were first isolated using Buffer 300 (50 mM Tris, pH7.5, 1% NP40, 1mM EDTA, 1mM $MgCl_2$, 300 nM NaCl). Lysates were subsequently incubated with 30 μL of Protein A/G beads coated with IgG, BRG1, or V5 antibodies overnight at 4°C (**Supplemental Table 4**). Beads were washed, and immunoprecipitates were separated on a gradient bis-tris gel (4-12%). BRG1, BAF155, SS18, and BAF47 were probed using corresponding antibodies (**Supplemental Table 4**). To detect BAF170, blots were stripped (LI-COR Biosciences) and probed with BAF170 antibody (**Supplemental Table 4**). Protein bands were visualized on the Odyssey CLx imager (LI-COR Biosciences).

##### Fluorescence Activated cell sorting analysis by flow cytometry

To detect changes in the protein expression of SDF2 and HBG, cells were first fixed and permeabilized in the Ctyofix/CytoPerm solution (BD Biosciences) according to manufacturer's protocol. Cells were washed twice with 1 x BD Perm/Wash buffer (BD Biosciences) and incubated with IgG, HBG, or SDF2 antibodies (**Supplemental Table 4**) for 1 h at 4 °C with the same buffer. Cells were then washed twice with 1x BD Perm/Wash buffer (BD Biosciences), and cells were stained with PE-conjugated anti-rabbit secondary antibody (ThermoFisher Scientific) (**Supplemental Table 4**). Antibody-stained cells were acquired on BD FACSymphony A3 Cell Analyzer (BD Biosciences), and data was analyzed using FlowJo software (BD).

##### Proliferation assay

Cell proliferation is assessed using CellTiter-Glo 2.0 Cell Viability Assay (Promega Corporation) according to manufacturer's protocol. In brief, an equal amount of CelTiter-Glo 2.0 is added to the cells and incubated in the dark for 2 mins while rotating at 700 x rpm. Cells were incubated at room temperature for an additional 10 mins, and luminescence values were obtained on the EnVision Multimode Plate Reader (PerkinElmer)

##### Statistical Analysis

When comparing 2 or more groups, statistical significance was determined by a one-way or two-way ANOVA with Dunnett's or Turkey's multiple comparisons (Prism 10, Graphpad).

**Supplemental Table 2. sgRNA sequences determined by the Broad sgRNA database and konermann *et al*[8].**

| Name | Gene/Location | sgRNA sequence |
|---|---|---|
| sg*HBG1* 1 | *HBG1/2* | GGCTAGGGATGAAGAATAAA |
| sg*HBG1* 2 | | CTTGACCAATAGCCTTGACA |
| sg*HBG1* 3 | | ATGCAAATATCTGTCTGAAA |
| sgHS1 1 | DNase I Hypersensitivity site 1 of LCR region | GTGGATATAGATAAGAGCTC |
| sgHS1 2 | | ACTATGCTGAGCTGTGATGA |
| sgHS1 3 | | GGGACTGAGAAGGCAATAGC |
| sgHS1 4 | | GACTCTCACTGCCTTTAGCT |
| sgHS2 1 | DNase I Hypersensitivity site 2 of LCR region | AATATGTCACATTCTGTCTC |
| sgHS2 2 | | GGACTATGGGAGGTCACTAA |
| sgHS2 3 | | GAAGGTTACACAGAACCAGA |
| sgHS2 4 | | GCCCTGTAAGCATCCTGCTG |
| sgHS3 1 | DNase I Hypersensitivity site 3 of LCR region | GAGAGATAGACCATGAGTAG |
| sgHS3 2 | | TGCCAGCCTATAACCCATCT |
| sgHS3 3 | | CCAGCTATCAGGGCCCAGAT |
| sgHS4 1 | DNase I Hypersensitivity site 4 of LCR region | AGTCCACCCCTTCTCGGCCC |
| sgHS4 2 | | AACCCTGCTCGGGAATGGGA |

| | | |
|---|---|---|
| sgHS4 3 | | GTCAGCCTAGTAGAGAGGCA |
| sg*BGLT3* 1 | *BGLT3* loci | AAATGACCTAATGCCCAGCA |
| sg*BGLT3 2* | | AATGACCTAATGCCCAGCAC |
| sg*BGLT3 3* | | CTTCACTGACAATATTCCAC |
| sg*BGLT3 4* | | CGTCTCAAACAAGCAATTTC |
| sgHBG-4kbp 1 | 4 kbp upstream of *HBG* | CATCAGCAGAGGCAGTCAGG |
| sgHBG-4kbp 2 | | ACCAGCTGCGAAACCAAGTA |
| sgHBG-4kbp 3 | | TCCCATGCAACTCAAAAGGT |
| sg*SDF2* 1 | *SDF2* | CAGACAGGTAAAGTTCCGAC |
| sg*SDF2* 2 | | GTACTATCTTTTCCCAGTCT |
| sg*SDF2* 3 | | GTGGGGGCTTAGCGCACTGG |
| sg*SPI1* 1 | *SPI1* | AGGGCTGGCCTGGGAAGCCA |
| sg*SPI1 2* | | GGGACTGAGAGGGATGACTT |
| sg*SPI1 3* | | GGTCCAGGCCCCCTGCCCAG |
| sgNT | Non-Targeting | CTGAAAAAGGAAGGAGTTGA |

**Supplemental Table 3. Primers used for ChIP-qPCR and ATAC-qPCR**

| Primer (location from *HBG1 TSS*) | | Sequence (5' - 3') | Application |
|---|---|---|---|
| *HBG1* qPCR set 2 (-1818 bp) | For | TCAGAGTTTTCCACATGCCC | ChIP, ATAC |
| | Rev | TGGCATTTGAAGTGGGTCCTT | |
| *HBG1* qPCR set 4 (-887 bp) | For | TCCACAGTACCTGCCAAAGAA | ChIP, ATAC |
| | Rev | CAGGGTTTTGGCGTAGCTCT | |
| *HBG1* qPCR set 6 (-473 bp) | For | CCCGTCAAAAATCCTGGACC | ChIP, ATAC |
| | Rev | TAGCTCCAGTGAGGCCTGTA | |
| *HBG1* qPCR set 5 (-164 bp) | For | GTCCCTGGCTAAACTCCACC | ChIP, ATAC |
| | Rev | AACTGCTGAAGGGTGCTTCC | |
| *HBG1* qPCR set 7 (+93 bp) | For | AGGAGAAACCCTGGGAAGGT | ChIP, ATAC |
| | Rev | GAAGCGACCTGGACTTTTGC | |
| *HBG1* qPCR set 8 (+566 bp) | For | AGCAGGGTGTGAGCTGTTTG | ChIP, ATAC |
| | Rev | ACATTACCACTGGGTCTCAGC | |
| *HBG1* qPCR set 10 (+1507 bp) | For | CTGAGCTCACTGCCCATGAT | ChIP, ATAC |
| | Rev | CCCTCAATATAAACCCTTTGTGGC | |
| h*GAPDH* | For | CATCTCAGTCGTTCCCAAAGT | ChIP |

| | | | |
|---|---|---|---|
| | Rev | TTCCCAGGACTGGACTGT | |
| ATAC qPCR control | For | CCAGGGTTGGCTCAGTATGAT | ATAC |
| | Rev | TAGAACCTCGCCCTGACACA | |

**Supplemental Table 4. Antibodies used for immunoblotting, Co-IP, ChIP, and Flow cytometry**

***Antibodies for immunoblotting***

| Primary Antibody | Clone/Cat# | Company | Dilution |
|---|---|---|---|
| Anti-HA Tag | C29F4/3742S | Cell Signaling Technologies | 1:1000 |
| Anti-SS18 | D6I4Z/ 21792S | Cell Signaling Technologies | 1:1000 |
| Anti-FKBP12 | ab2918 | Abcam | 1:1000 |
| Anti-GAPDH | 6C5/ sc-32233 | Santa Cruz Biotechnology | 1:2000 |
| Anti-HBG | EPR9708(B)/ab137096 | Abcam | 1:1000 |
| Anti-BAF155 | D8I3U/94962S | Cell Signaling Technologies | 1:1000 |
| Anti-BRG1 | EPNCIR111A/ab110641 | Abcam | 1:5000 |
| Anti-BAF47(Ini1) | A5/sc-166165 | Santa Cruz Biotechnology | 1:250 |
| Anti-BAF170 | D8O9V/12760S | Cell Signaling Technologies | 1:2000 |
| Anti-SPI1 | EPR25123-110/ab302623 | Abcam | 1:1000 |

| Secondary Antibody | Clone/Cat# | Company | Dilution |
|---|---|---|---|
| IRDye 680RD Donkey anti-Mouse IgG | 926-68072 | Li-COR Biosciences | 1:10000 |

| | | | |
|---|---|---|---|
| IRDye 800CW Donkey anti-Rabbit IgG | 926-32213 | Li-COR Biosciences | 1:10000 |

***Antibodies for immunoprecipitation and co-immunoprecipitation***

| **Antibodies** | **Clone/Cat#** | **Company** | **Amount** |
|---|---|---|---|
| Anti-HBG | EPR23381-254/ab283313 | Abcam | 3 µg |
| Anti-IgG | sc-2025 | Santa Cruz Biotechnology | 3 µg |
| Anti-V5 | SV5-Pk1/ab27671 | Abcam | 3 µg |
| Anti-BRG1 | G-7/sc-17796 | Santa Cruz Biotechnology | 4 µg |

***Antibodies for ChIP***

| **Antibodies** | **Clone/Cat#** | **Company** | **Amount** |
|---|---|---|---|
| Anti-V5 | SV5-Pk1/R-960-25 | ThermoFisher Scientific | 5 µg |
| Anti-BRG1 | 13-2002 | Epicypher | 5 µl |
| Anti-IgG | DA1E/3900S | Cell Signaling Technologies | 5 µg |
| Anti-H3 | 39763 | Active Motif | 5 µg |
| Anti-H3K4me3 | 15-10C-E4/05-745R | Millipore Sigma | 3 µl |
| Anti-H3K27Ac | EP16602/ab177178 | Abcam | 5 µg |

| | | | |
|---|---|---|---|
| Anti-H3K27me3 | 39155 | Active Motif | 5 µg |
| Anti-GLTSCR1 | E6I3A/45441S | Cell Signaling Technologies | 10 µL |
| Anti-ARID1A | D2A8U/12354S | Cell Signaling Technologies | 5 µL |
| Anti-ARID2 | D8D8U/82342S | Cell Signaling Technologies | 20 µL |
| Anti-RING1B | D22F2/5694S | Cell Signaling Technologies | 10 µL |
| Anti-H2AK119Ub1 | D27C4/8240S | Cell Signaling Technologies | 5 µL |
| Anti-EZH2 | AC22/39875 | Active Motif | 3 µg |

***Antibodies for flow cytometry***

| **Primary Antibody** | **Clone/Cat#** | **Company** | **Dilution** |
|---|---|---|---|
| Anti-IgG | 2729S | Cell Signaling Technologies | 1:100 |
| Anti-HBG | 25728-1-AP-150UL | Proteintech | 1:100 |
| Anti-SDF2 | 14747-1-AP | Proteintech | 1:100 |

| **Secondary Antibody** | **Clone/Cat#** | **Company** | **Dilution** |
|---|---|---|---|
| Goat anti-Rabbit IgG (H+L) Cross-Adsorbed Secondary Antibody, PE | P-2771MP | ThermoFisher Scientific | 1:500 |

### Uncut Blot

#### Figure 2B: Uncropped gels

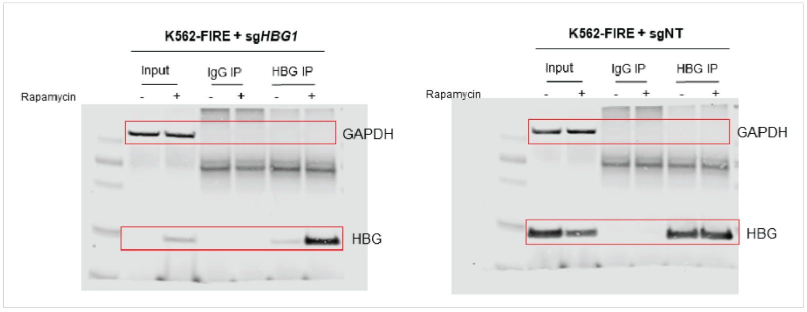


IP: 3 ug Ab (ab283313)
Lysate: 0.75 mg Proteins
Input: 30 ug
Primary Ab: HBG (1:1000, ab137096)
GAPDH (1:2000, sc-32233)
Secondary AB: anti ms (1:10000, 926-68072)
anti rb (1:10000, 926-32213)

#### Figure 5F: Uncropped gels

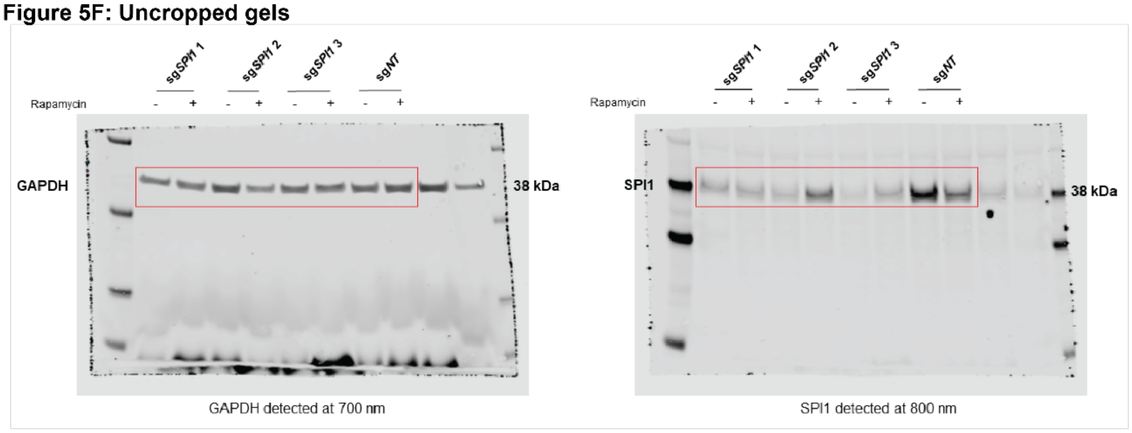


Sample: 30 ug protein
Primary Ab: SPI1 (1:1000, ab302623, Abcam)
GAPDH (1:2000, sc-32233, Santa Cruz)
Secondary AB: anti ms (1:10000, 926-68072)
anti rb (1:10000, 926-32213)

#### Supplemental Figure 1B: Uncropped gels

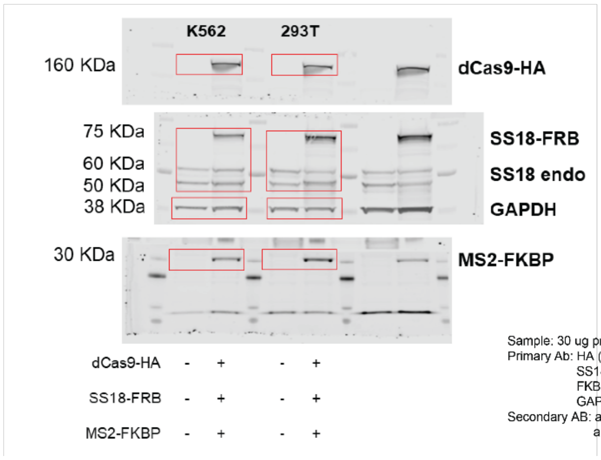


Sample: 30 ug protein
Primary Ab: HA (1:1000, 3724S, CST)
SS18 (1:1000, 21792S, CST)
FKBP (1:1000, ab2918, Abcam)
GAPDH (1:2000, sc-32233, Santa Cruz)
Secondary AB: anti ms (1:10000, 926-68072)
anti rb (1:10000, 926-32213)

#### Supplemental Figure 1C: Uncropped gels

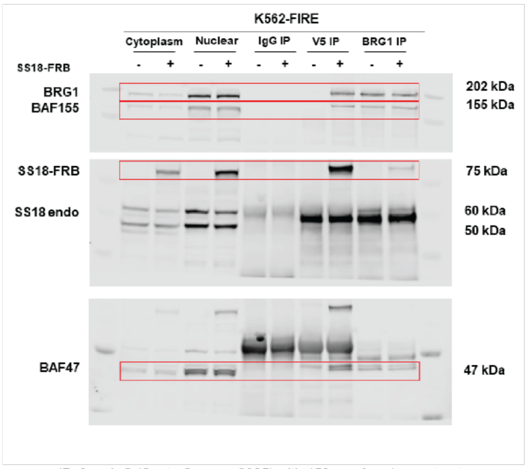


IP: 3 ug IgG (Santa Cruz, sc-2025) with 450 ug of nuclear extract
3 ug V5 (Abcam, ab27671) 450 ug of nuclear extract
4 ug BRG1 (Santa Cruz, sc-17796) with 1 mg of nuclear extract
Cyto and nuclear Input: 30 ug
Primary Ab: BAF155 1:1000 (CST, 94962)
BRG1 1:5000 (Abcam 110641)
SS18 1:1000 (CST, 21792S)
BAF47 1:250 (Santa Cruz, sc-166165)
Secondary AB: anti ms (1:10000, 926-68072)
anti rb (1:10000, 926-32213)

#### Supplemental Figure 1D: Uncropped gels

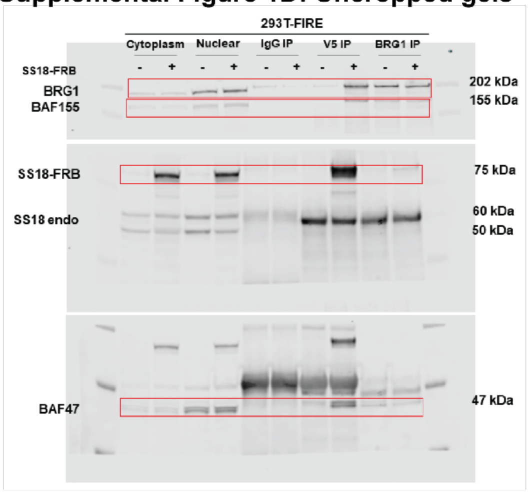


IP: 3 ug IgG (Santa Cruz, sc-2025) with 450 ug of nuclear extract
3 ug V5 (Abcam, ab27671) 450 ug of nuclear extract
4 ug BRG1 (Santa Cruz, sc-17796) with 1 mg of nuclear extract
Cyto and nuclear Input: 30 ug
Primary Ab: BAF155 1:1000 (CST, 94962)
BRG1 1:5000 (Abcam 110641)
SS18 1:1000 (CST, 21792S)
BAF47 1:250 (Santa Cruz, sc-166165)
Secondary AB: anti ms (1:10000, 926-68072)
anti rb (1:10000, 926-32213)

#### Supplemental Figure 1C-D: Uncropped gel for BAF170

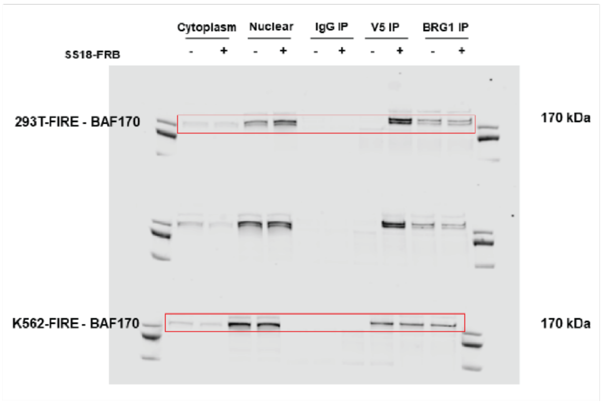


Blot initially stained with BAF155 and BRG1 was stripped with 1x stripping buffer (LiCor) and reblotted o/n with BAF170 antibody

IP: 3 ug IgG (Santa Cruz, sc-2025) with 450 ug of nuclear extract
3 ug V5 (Abcam, ab27671) 450 ug of nuclear extract
4 ug BRG1 (Santa Cruz, sc-17796) with 1 mg of nuclear extract
Cyto and nuclear Input: 30 ug
Primary Ab: BAF170, 1:2000 (CST, 12760S)
Secondary AB: anti ms (1:10000, 926-68072)
anti rb (1:10000, 926-32213)
